# Receptor-Like Kinase Phosphorylation of Arabidopsis Heterotrimeric G-Protein Gα - Subunit AtGPA1

**DOI:** 10.1101/752337

**Authors:** Haiyan Jia, Gaoyuan Song, Emily G. Werth, Justin W. Walley, Leslie M. Hicks, Alan M. Jones

## Abstract

As molecular on-off switches, heterotrimeric G protein complexes, comprised of a Gα subunit and an obligate Gβγ dimer, transmit extracellular signals received by G protein– coupled receptors (GPCRs) to cytoplasmic targets that respond to biotic and abiotic stimuli. Signal transduction is modulated by phosphorylation of GPCRs and G protein complexes. In *Arabidopsis thaliana*, the Gα subunit AtGPA1 is phosphorylated by the receptor-like kinase (RLK) BRI1-ASSOCIATED Kinase 1 (BAK1), but the extent that other RLKs phosphorylates AtGPA1 is unknown. We mapped 22 trans-phosphorylation sites on AtGPA1 by 12 RLKs hypothesized to act in the Arabidopsis G protein signaling pathway. Cis-phosphorylation sites on these RLKs were also identified. BRI1, BAK1, and SERK1 have been reported as Ser/Thr and Tyr dual specificity kinases. We identified 4 more dual specificity kinases: IOS1, PSY1R, PEPR1, and AT2G37050. Multiple sites are present in the core AtGPA1 functional units, including pSer52 and pThr53 of the conserved P-loop that directly binds nucleotide/phosphate, pThr164 and pSer175 from αE helix in the intramolecular domain interface for nucleotide exchange and GTP hydrolysis, and pThr193 or pThr194 in Switch I (SwI) that coordinates nucleotide exchange and protein partner binding. Several AtGPA1 S/T phosphorylation sites are nucleotide-dependent phosphorylation patterns, such as S52/T53 in the P-loop and T193 and/or T194 in SwI.

---

Heterotrimeric G protein complexes link extracellular signals perceived by G protein– coupled receptors (GPCRs) to downstream effectors in regulating cellular responses^[1]^. The G protein complex is composed of a Gα subunit that binds GDP and GTP, a Gβ and Gγ dimer wherein a cycle of GTP binding activates this complex and GTP hydrolysis deactivates.^[2]^ GTP binding is catalyzed by GPCRs in animals but is spontaneous in plants.^[3]^ This GTPase cycle is modulated by several proteins, such as GTPase-accelerating proteins (GAPs; e.g. regulator of G signaling (RGS) proteins^[4]^), GDP-dissociation inhibitors (GDI), and receptor and nonreceptor guanine-nucleotide exchange factors (GEFs; e.g. GPCRs and GIV).

Phosphorylation of 7 transmembrane (7TM) GPCRs is a regulatory mechanism to modulate G protein signaling.^[5]^ In animals, extracellular signals induce GPCR phosphorylation by G protein–coupled receptor kinases (GRKs) on the C-terminus and intracellular loops leading to arrestin recruitment and signal desensitization through receptor endocytosis. Phosphorylation patterns of GPCR are recognized by arrestins via a phospho-barcoding mechanism to activate arrestin-dependent effectors in diverse cellular processes.^[5]^ In *Arabidopsis thaliana*, AtGPA1 is kept in its deactivated state by a 7TM GAP (AtRGS1) and the ligand-induced phosphorylation and endocytosis of AtRGS1 activates AtGPA1 via de-repression.^[6]^

In animals, a few Gα phosphosites have known functions in G signaling.^[7]^ N terminal pSer16 (phosphorylated by PKA or PKC) and pSer27 (phosphorylated by PKC) in Gα_z_ prevent its binding to Gβγ (pSer16) and RGS.^[8, 9]^ SRC phosphorylation of Y37 and Y391 in Gα_s_ increase receptor-stimulated GTPγS-binding and GTP hydrolysis.^[10, 11]^ YpkA phosphorylation of Gα_q_ Ser53 in the P-loop impairs GTP-binding and Gα activation.^[12]^ Some of these phosphorylated residues plus pY166 are conserved in animals and plants, and the crystal structure of AtGPA1 (PDB 2XTZ) is highly similar (RMSD = 1.8 ◻) to animal G subunits.^[3]^ *In vivo* AtGPA1 phosphorylation at Y166 is induced by hormones.^[13]^ A mechanism termed “substrate phosphoswitching” proposes that phosphorylation of pY166 switches AtRGS1 from a GAP to a quasi GDI.^[14]^ The AtGPA1^Y166E^ phosphomimetic mutant reduces the AtRGS1-accelerated GTP hydrolysis rate.^[14]^

Although SAPH-ire (Structural Analysis of PTM Hot spots^[15]^) predicted multiple key modified alignment positions in the Gα family, including N-terminal α-helix (αN), P-loop, and αE helix,^[14]^ the presence of those predicted phosposites needs to be verified experimentally. AtGPA1 was shown as substrate of the S/T and Y dual-specificity RLK, BAK1.^[14, 16]^ Arabidopsis has approximately 400 RLKS.^[17]^ A set of 70 LRR RLKs was screened for AtGPA1 phosphorylation and 18 RLKs were identified as AtGPA1 phosphokinases.^[14]^ Recently, Xue et al^[18]^ reported that BAK1 was necessary for flg22– induced AtGPA1 phosphorylation *in vivo*. They further identified 16 BAK1 mediated AtGPA1 phosphosites and determined that T19 was necessary for BAK1 *in vitro* phosphorylation of AtGPA1.

Here, we explore the RLK-mediated phosphorylation of AtGPA1 as a mechanism for regulation of G protein signaling in plants. Except partially for BAK1,^[18]^ RLK-mediated AtGPA1 phosphorylation sites have not been exhaustively identified.

Twelve RLKs previously shown to phosphorylate AtGPA1^[14]^ were chosen for this study (**Table 1**) to identify AtGPA1 phosphorylation sites via LC-MS-MS (See Supporting Information Methods for details). Briefly, 5 µg of each RLK was mixed with 15 μg twinstrep or His-tagged AtGPA1 in kinase reaction buffer in presence of 50 µM GDP or 100 µM GTPγS. The kinase reaction samples were used for proteomic analysis to detect phosphorylated residues (hereafter P-sites) of AtGPA1 and the corresponding RLKs. Products of *in vitro* kinase reactions were trypsin digested. For reactions of RLKs with twinstrep-AtGPA1, tryptic peptides were subjected to LC-MS-MS analysis using a Q-Exactive Plus high-resolution quadrupole Orbitrap mass spectrometer.^[19]^ Maxquant^[20]^ was used to identify peptides and locate the phosphorylation sites (**Figure 1; Table S1-S2**). For the BAK1 reaction with his-AtGPA1 (**Table S3 and Figure S2**), NanoAcquity UPLC (Waters) and TripleTOF 5600 (AB Sciex, https://sciex.com/) mass spectrometer were used for LC-MS-MS analysis. Peptide sequence determination and protein inference were done by Mascot (v2.5.1; Matrix Science) using the TAIR website (https://www.arabidopsis.org/download/index-auto.jsp?dir=%2Fdownload_files%2FProteins%2FTAIR10_protein_lists).

**Table 1.**
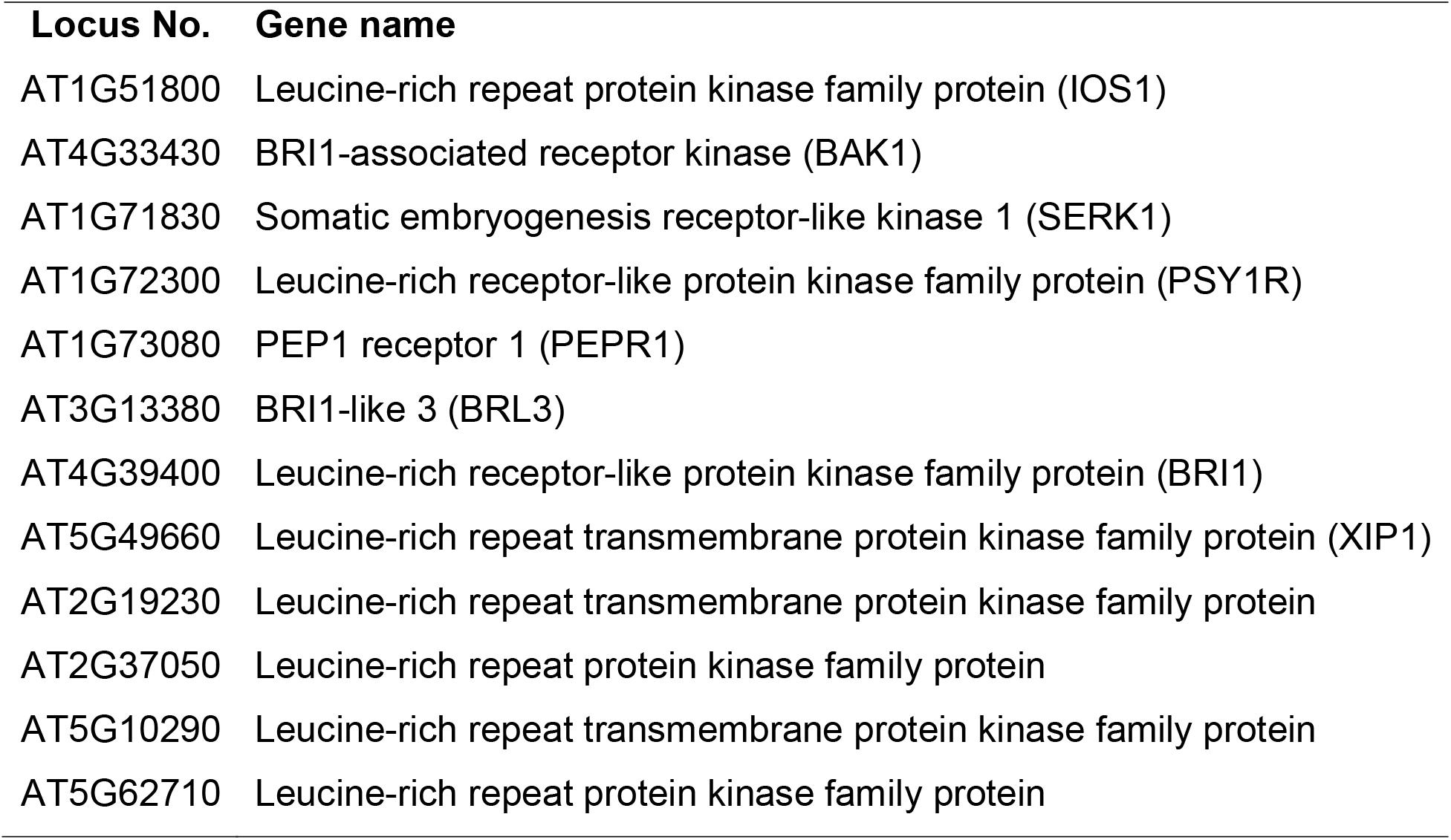
AtGPA1 RLKs used in this study.

| **Locus No.** | **Gene name** |
| --- | --- |
| AT1G51800 | Leucine-rich repeat protein kinase family protein (IOS1) |
| AT4G33430 | BRI1-associated receptor kinase (BAK1) |
| AT1G71830 | Somatic embryogenesis receptor-like kinase 1 (SERK1) |
| AT1G72300 | Leucine-rich receptor-like protein kinase family protein (PSY1R) |
| AT1G73080 | PEP1 receptor 1 (PEPR1) |
| AT3G13380 | BRI1-like 3 (BRL3) |
| AT4G39400 | Leucine-rich receptor-like protein kinase family protein (BRI1) |
| AT5G49660 | Leucine-rich repeat transmembrane protein kinase family protein (XIP1) |
| AT2G19230 | Leucine-rich repeat transmembrane protein kinase family protein |
| AT2G37050 | Leucine-rich repeat protein kinase family protein |
| AT5G10290 | Leucine-rich repeat transmembrane protein kinase family protein |
| AT5G62710 | Leucine-rich repeat protein kinase family protein |

**Figure 1.**
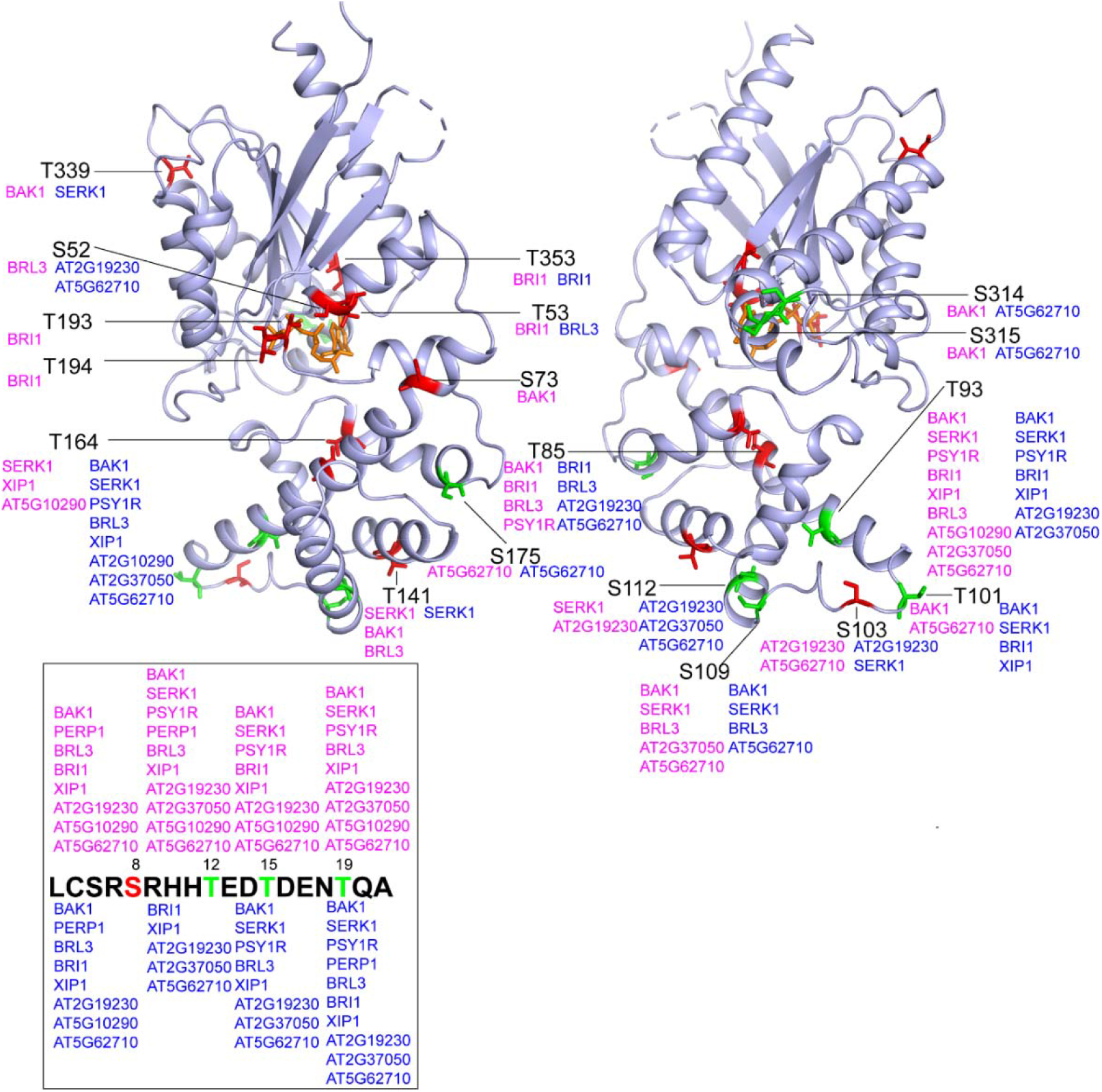
RLK phosphorylation sites on GPA1 (GDP) and GPA1 (GTP) states identified by LC-MS/MS. Multiple RLK P-sites on AtGPA1 were detected by LC-MS/MS. To show the P-sites in the front and back view, the AtGPA1 structure was rotated 180°C. S/T residues that are conserved in animals and plants are labelled in red. S/T residues that are specific in plants are labelled in green. The positions of corresponding AtGPA1 S/T residues are labelled by solid black lines. RLKs that catalyze the phosphorylation reaction in the (AtGPA1•GDP) state and (GPA1•GTP) state are shown in magenta and blue, respectively. For orientation, S52/T53 residues are in the P-loop, and T193/T194 residues are in Switch I. Phosphorylation patterns of S/T residues located at the GPA1 N terminus (missing in the structure) are shown in the box.

We identified AtGPA1 P-sites from all RLKs except for IOS1 (**Figure 1 and Table S1**) despite multiple IOS1 autophosphorylation sites suggesting that this recombinant IOS1 is an active kinase that does not recognize AtGPA1 as a substrate (**Table S2**). We speculate that the previously strong phosphorylation intensity detected by autoradiography^[14]^ was misinterpreted due to a similar-size phosphorylated IOS1 or an unknown co-purified protein. All other tested RLKs had detected autophosphorylation sites (**Table S2**).

We found that 11 of the tested RLKs phosphorylate AtGPA1 at 22 Ser/Thr residues (**Figure 1, S2, Table S1 and S3**) of which 11 are conserved in both animal and plant Gα subunits and the rest are plant Gα subunit specific (**Figure S1**). Twelve P-sites were found in the RAS-like domain where P-loop and Switch I to III (SwI to III) are localized (**Figure S1**).^[3]^ Ten P-sites were present in the all-helical domain which is essential for AtGPA1 rapid nucleotide exchange and self-activation properties.^[3, 21]^ Molecular dynamic simulation illustrates that rapid nucleotide exchange in AtGPA1 is partly determined by intradomain interaction in the all-helical domain, especially helices αA and αB (**Figure S1**).^[21]^ Therefore, P-sites localized at αA (pSer73, pThr85 and pThr93) and αB (pSer109 and pSer112) helices, and the linker (pSer101 and pSer103) between αA-αB, potentially regulate nucleotide exchange and AtGPA1 self-activation (**Figure 1 and S1**). SAPH-ire predicted three key MAP clusters in the Gα family, including the N-terminal α-helix (αN), P-loop, and αE helix.^[14]^ We validated this prediction with identification of 4 P-sites in αN, which included pSer8, pThr12, pThr15 and pThr19, that may regulate AtRGS1 and Gβγ binding, given that the αN is interfaces with receptors^[22]^ and Gβγ.^[23, 24]^ pThr12, pThr15 and pThr19 were reported as in vivo AtGPA1phosphosites.^[13, 25–27]^ Ser52 and Thr53 are present in the conserved P-loop that directly binds nucleotide/phosphate, suggesting pSer52 and pThr53 affect nucleotide binding and/or GTPase activity. Consistent with mutational analyses showing that phosphorylation of this Ser in Gα_q_ impairs GTP-binding.^[12]^ Two P-sites were detected on helix αE, specifically, pThr164 and pSer175. Like Tyr166,^[14]^ residues Thr164 and Ser175 on AtGPA1 are positioned in the intramolecular domain interface where nucleotide exchange and GTP hydrolysis occurs (**Figure 1**),^[3]^ suggesting that pThr164 and pSer175 may control GPA1 GTP binding and/or GTPase activity (intrinsic and/or accelerated by AtRGS1). While pTyr166 is predicted to be a conserved PTMs on helix αE^[14]^ and is detected *in vivo* from mass spec analysis of phytohormone treated Arabidopsis cell cultures,^[13]^ no pTyr166 peptides were detected. This could be caused by non-optimal *in vitro* phosphorylation conditions, technical issues of mass spec analysis and/or a prerequisite PTM that occurs *in vivo*. We also identified 2 P-sites, pThr193 and pThr194, in SwI (**Figure 1 and S1**), where nucleotide-induced AtGPA1 conformational changes occur.^[28]^ T193 and T194 are likely residues contacting AtRGS1 and Gβγ.^[29]^ As such, we propose that pThr193 and pThr194 modulate AtGPA1 nucleotide exchange and AtRGS1/Gβγ binding. Additional P-sites, such as pSer314 and/or pSer315, pThr339, and pThr353, localized at the C-terminal part of the RAS domain distal to the P-loop and switches, may regulate AtGPA1 effector interactions.

Nucleotide-dependent phosphorylation was observed for multiple AtGPA1 Ser/Thr sites, including Ser52, Thr53, Thr193, Thr194, Ser73, Thr339, Ser314 and Ser315 (**Figure 1**). For example, Ser52 was only phosphorylated by BRL3 in the AtGPA1•GDP state, and was phosphorylated by AT2G19230 and AT5G62710 in the AtGPA1•GTP state. While T53 was BRI1-phosphorylated in the AtGPA1•GDP state and is BRl3-phosphorylated in the AtGPA1•GTP state. This reveals that nucleotide-induced conformation change in AtGPA1 alters RLK’s specificity possibly by altering accessibility to the S52 and T53 substrates; Two important residues in the AtGPA1 SwI, T193 and T194, were phosphorylated by BRI1 only in the AtGPA1•GDP state. Residues T193 and T194 when in the AtGPA1•GTP state were not phosphorylated by any of the tested RLKs suggesting that T193 and 194 are buried in the GTP-bound state. Additionally, only BAK1 phosphorylates S73 in the AtGPA1•GDP state. BAK1 and AT5G62710 phosphorylate S314 and S315 in the GDP-bound state and GTP-bound state, respectively. Residue T339 is a substrate of BAK1 in its GDP-bound state and a substrate of SERK1 in the GTP-bound state. In depth comprehensive investigations are on-going to illustrate the functional significance of these nucleotide-dependent AtGPA1 P-sites.

Twelve Ser/Thr sites were RLK-phosphorylation hot spots (defined as phosphorylated by more than 3 tested RLKs) (**Figure 1**). They are Ser8, Thr12, Thr15, Thr19, Thr85, Thr93, Thr101, Ser103, Ser109, Ser112, Thr141 and Thr164. This suggest that phosphorylation of these sites by RLKs is necessary for AtGPA1 mediated physiological responses.

A database of auto phosphorylation sites of 73 LRR RLKs was reported.^[30]^ SERK1,^[31]^ BRI1 and BAK1^[16]^ are Ser/Thr and Tyr dual specificity kinases among the 12 tested kinases. We identified 4 more dual specificity kinases (**Table S2**): IOS1, PSY1R, PEPR1, and AT2G37050. Cis phosphorylation of IOS1 is known to occur at 12 S/T sites^[30]^ and we show here that IOS1 auto phosphorylated at 32 sites including pY697. We detected 19 autophosphorylation sites from PSY1R, including two tyrosine P-sites Y837 and Y865. Four tyrosine P-sites (pY805,831,901,and 910) were observed among total 16 P-sites of PEPR1. Twenty-one auto P-sites were identified from AT2G37050 including pY717.

## Data Availability

Raw data files and MaxQuant Search results were deposited in the Mass Spectrometry Interactive Virtual Environment (MassIVE) repository: https://massive.ucsd.edu/ProteoSAFe/static/massive.jsp with dataset identifier: MSV000083963 (Kinase reactions with GDP) and MSV000083964 (Kinase reactions with GTP).” Raw (*.wiff) data files from the TripleTOF 5600 and peak lists (*.mgf) were deposited in the Mass Spectrometry Interactive Virtual Environment (MassIVE) repository with dataset identifier: MSV000084139 (His-GPA1 Kinase reaction with GDP-Hicks).

## Acknowledgements

This work was supported by NIH (1R01GM120316-01A1), NSF (1759023), and by the ISU Plant Sciences Institute awarded to Justin W. Walley and NIGMS (R01GM065989), NSF (MCB-1552522) awarded to Leslie M. Hicks, NIGMS (R01GM065989) and NSF (MCB-0718202) awarded to Alan. M. Jones.

