## Supporting Information for "Receptor-Like Kinase Phosphorylation of Arabidopsis Heterotrimeric G-Protein Gα - Subunit AtGPA1"

**Protein expression and purification**

Twinstrep AtGPA1 was cloned into the pDEST17 vector (His tag removed) and transformed into ArcticExpress RP cells (Agilent Technologies). At OD600 0.6-0.8, protein expression was induced using 0.5 mM IPTG and cultured at 12 °C for 16 hours. All purification steps were performed at 4 °C. Cell pellets were resuspended in extraction buffer (25 mM Tris-HCl pH 8.0, 150 mM NaCl, 2 mM MgCl2, 20 µM GDP, 5 mM 2-mercaptoethanol, 1 mM PMSF, 0.25 mg/mL Lysozyme, 0.1% Thesit (Sigma, 88315), 1 X protease inhibitor cocktail, 10% glycerol) and mixed for 30 min. The suspension was sonicated (Sonic Dismembrator, Model 550, Fisher Scientific, power level 5, 0.50/0.50 off for 1 min, 2 cycles). The soluble fraction was obtained by centrifugation at 30,000 X g for 40 mins, then the soluble fraction was incubated 30 mins with strep-tactin sepharose (50% suspension, cat no. 2-1201-010, IBA). The resin was washed with washing buffer (25 mM Tris-HCl pH 8.0, 150 mM NaCl, 2 mM MgCl2, 20 µM GDP, 5 mM 2-mercaptoethanol, 1 mM PMSF, 1 X protease inhibitor cocktail ,10% glycerol) and eluted with elution buffer (25 mM Tris-HCl pH 8.0, 150 mM NaCl, 2 mM MgCl2, 20 µM GDP, 5 mM 2-mercaptoethanol, 1 mM PMSF, 1 X protease inhibitor cocktail, 10% glycerol, 2.5 mM desthibiotin (Sigma-Aldrich)). The monodispersed fraction was obtained by loading the eluate on size exclusion column (Superdex 200 10/300 GL, GE Healthcare) with running buffer (20 mM Tris-HCl, pH 7.5, 50 mM NaCl, 10 mM MgCl2, 50 µM GDP, 1 mM DTT, and 10% Glycerol). Aliquoted protein samples were snap frozen by liquid nitrogen and store at -80 °C.

RLKs were expressed and purified as previously described.^[1]^ RLKs-cDNA encoding the complete cytoplasmic domain was cloned into pET15b vector and transformed into BL21 (DE3) cells. Protein expression was induced using 0.5mM IPTG at OD600 0.6-0.8 at 30 °C for 3 hours. RLKs were purified by Ni-NTA Agarose (Qiagen, Mat. No, 1018244) and eluted with elution buffer (50 mM Tris-HCl, pH 8.0, 300 mM NaCl, 1 mM MgCl2, 250 mM imidazole, 5 mM 2-mercaptoethanol, 1 mM phenylmethyl-sulfonyl fluoride (PMSF)). And then the eluate was dialyzed against dialysis buffer (10 mM Tris-HCl, pH 8.0, 20 mM NaCl, 1 mM DTT, 1 mM PMSF, 10% glycerol), 4 °C overnight. Aliquoted protein samples were snap frozen by liquid nitrogen and store at -80 °C.

**In vitro kinase assays**

Purified 5 µg of each RLK was mixed with 15 μg of Twinstrep-AtGPA1 in kinase reaction buffer to 30ul reaction (50 mM Tris-HCl, pH 7.5, 10 mM MgCl2, 10 mM MnCl2, 2 mM Na3VO4, 1 mM DTT, 1 μg/ml leupeptin, 0.1 μM calyculin A, 50 μM ATP, 50 μM GDP or 100 μM GTPγS). Control reactions have AtGPA1 only. The above reaction was incubated at 25°C for 4 hours. And then the samples were snap frozen by liquid nitrogen and store at -80 °C. Additionally, same kinase reaction condition was performed for 5ug of BAK1 and 20ug of His-GPA1 in presence of GDP. The reaction products were separated by SDS-PAGE gel and stained with Coomassie Blue.

**LC-MS/MS and phosphorylation site detection**

The products of in vitro kinase reactions of twinstrep-GPA1 were processed into peptides and analyzed by LC-MS/MS based on established methods as follows.^[2-4]^ Proteins (~50μl) were mixed with 400 μl of 8 M urea in 0.1 M Tris-HCl pH 8.0 (UA buffer) and 1 x phosphatase inhibitor (2.5 mM sodium fluoride, 0.25mM sodium vanadate, 0.25mM sodium pyrophosphate decahydrate, and 0.25mM glycerophosphate in H_2_O). The samples were then added to a Microcon-30kDa centrifugal filter (Cat # MRCF0R030) and centrifuged at 12,000 x g for 15 min. To UA buffer (200 μl),1 x phosphatase inhibitor was added to the filtrate and centrifuged at 12,000 x g for 10 min. UA buffer supplemented with 200 μl 2 mM Tris(2-carboxyethyl)phosphine hydrochloride (TCEP-HCl, Thermo Scientific) and 1 x phosphatase inhibitor was added to the filtrate, then centrifuged. UA buffer supplemented with 50 mM iodoacetamide (IAM) with 1 x phosphatase inhibitor was added to the filtrate, which was incubated in the dark for 30 min and then centrifuged. The samples were then processed 4 times with 200 μl UA buffer with 1 x phosphatase inhibitor and twice with 50 mM ammonium bicarbonate with 1 x phosphatase inhibitor. Proteins were then digested with 1μg trypsin (Roche, Cat# 03708969001) in 50 mM ammonium bicarbonate with 1 x phosphatase inhibitor overnight at 37°C, then digested with 0.5μg trypsin and 0.05 μg Lys-C at 37°C for 3 hours. Recovered peptides were acidified to a pH of ~2-3 with formic acid and desalted with 50 mg Sep-Pak C18 cartridges (Waters). Eluted peptides were dried using a SpeedVac and resuspended in 0.1% formic acid. Peptide amount was then quantified using the Pierce BCA Protein assay kit (Cat # 23225). Peptides (0.8 μg) of were loaded onto a 5 cm capillary column packed with 5 μM Zorbax SB-C18 (Agilent), which was connected via a zero dead volume 1 μm filter (Upchurch, M548) to a 20 cm nanospray tip packed with 2.5 μM C18 (Waters).

Peptides were separated using an acetonitrile gradient of 5-30% for 120 min, 30-80% for 25 min, and 0% for 5 min (150 min total) that was delivered via an Agilent 1260 quaternary HPLC at a flow rate of ~500 nL min-1. The HPLC system was coupled with a Thermo Scientific Q-Exactive Plus high-resolution quadrupole Orbitrap mass spectrometer using a custom fabricated nano-spray source. Data dependent acquisition was obtained using Xcalibur 4.0 software in positive ion mode with a spray voltage of 2.00 kV, and RF of 60, and a capillary temperature of 275°C. MS1 spectra were measured at a resolution of 70,000 with an automatic gain control (AGC) of 3e6, a maximum ion time of 100 ms, and a mass range of 400-2000 m/z. Up to 15 MS2, with a charge state of 2 to 4, were triggered at a resolution of 17,500, an AGC of 1e3 with a maximum ion time of 50 ms, a 1.5 m/z isolation window, and a normalized collision energy of 28. MS1 that triggered MS2 scans were dynamically excluded for 25 seconds.

**Phosphopeptide enrichment of His-GPA1 (GDP)**

For BAK1 and His-GPA1 sample, Coomassie Blue stained band at 45 kDa containing His-GPA1 protein was excised manually for downstream processing. Gel slices were de-stained and processed as previously described.^[5]^ Briefly, the gel band was de-stained using 50 mM ammonium bicarbonate/50% acetonitrile (MeCN) solution, reduced with 10 mM dithiotreitol (30 min, RT), alkylated with 55 mM iodoacetamide (30 min, RT, dark) and an in-gel trypsin digestion (25 ng trypsin in 50 mM ammonium bicarbonate) was preformed overnight at 37C as previously described.^[6]^ Peptides were extracted with 1% formic acid in 2% MeCN followed by a second extraction with 60% MeCN. Peptide extracts were dried by vacuum centrifugation.

Following peptide extraction from SDS-PAGE gel, peptides were subjected to a phosphopeptide enrichment using a 1 mg Titansphere Phos-TiO2 Kit spin columns (GL Sciences, http://www.glsciences.com/). Peptides were resuspended in 80% MeCN, 1% trifluoroacetic acid (TFA) and the spin column was conditioned twice with 80% MeCN, 1% TFA followed by centrifugation at 1000 g for 2 min. After this, peptides were loaded onto the spin column, centrifuged at 1000 g for 5 min, and flow-through was collected for LC-MS/MS analysis (“unenriched”). Following this, the column was washed four times using 80% MeCN, 1% TFA, and then phosphopeptides were eluted from the enrichment column with 20% MeCN, 5% ammonium hydroxide (“enriched”). After enrichment, both the flow-through and the eluent from the spin column were dried down using vacuum centrifugation and subjected to LC-MS/MS analysis.

**LC-MS/MS and phosphorylation site detection of His-GPA1 (GDP)**

Peptides were resuspended in 95% H2O/5% MeCN/0.1% TFA before separation via a 30 min linear gradient from 95% H2O/5% MeCN/0.1% formic acid (FA) to 60% H2O/40% MeCN/0.1% FA via a NanoAcquity UPLC (Waters). A TripleTOF 5600 (AB Sciex, https://sciex.com/) mass spectrometer was operated in positive-ionization and high-sensitivity mode for data acquisition as previously described,^[7]^ with the first 20 features above 150 counts threshold having a charge state of +2 to +5 selected for fragmentation during every 2 sec cycle.

**Data analysis**

For RLK reactions with twinstrep-GPA1, the raw data were analyzed using MaxQuant version 1.6.3.3.^[8]^ Spectra were searched, using the Andromeda search engine,^[9]^ against the Tair10 proteome file entitled “TAIR10_pep_20101214” that was downloaded from the TAIR website (https://www.arabidopsis.org/download/index-auto.jsp?dir=%2Fdownload_files%2FProteins%2FTAIR10_protein_lists) and was complemented with reverse decoy sequences and common contaminants by MaxQuant. Carbamidomethyl cysteine was set as a fixed modification while methionine oxidation, protein N-terminal acetylation, and protein phosphorylation (SYT) were set as variable modifications. The digestion parameters were set to “specific” for “Trypsin/P;LysC” with a maximum of 2 missed cleavages. The match between runs feature was turned off. Default settings were used for the remaining parameters including a peptide spectrum match and protein false discovery rate (FDR) of 0.01, which was determined using a reverse decoy database.

For BAK1 reaction with His-GPA1, Peptide sequence determination and protein inference was done by Mascot (v2.5.1; Matrix Science) using the TAIR website (https://www.arabidopsis.org/download/index-auto.jsp?dir=%2Fdownload_files%2F Proteins%2FTAIR10_protein_lists) appended with sequences for common laboratory contaminants (http://thegpm.org/cRAP/; 116 entries). For database searching, trypsin protease specificity with up to two missed cleavages, peptide/fragment mass tolerances of 50 ppm/0.1 Da, a fixed modification of carbamidomethylation at cysteine, and variable modifications of acetylation at the protein N-terminus, oxidation at methionine, deamidation at asparagine or glutamine, phosphorylation at serine (Ser) or threonine (Thr) and phosphorylation at tyrosine (Tyr) were used. Peptide false discovery rates (FDR) were adjusted to ≤1%. Based on Mascot Delta scoring^[10]^ for site-localization, T15, T19, S73, S112 were localized with >99% confidence, S109 with >95% confidence, and S314 or S315 could not be distinguished (50% probability for both).
